# LysinFusion: Integrating Multi-Feature Encoding and Hybrid CNN-Transformer Architecture for Phage Lysin Prediction

**DOI:** 10.64898/2026.03.13.711743

**Authors:** Sinuo He, Hanshuo Lu, Zhaomin Yao, Hanjun Zhang, Yihuai Cai, Fengfeng Zhou, Xin Feng, Yueyi Cai, Fei Li

## Abstract

The escalating antimicrobial resistance crisis has intensified the urgent need for alternative antibacterial agents. Phage lysins exhibit potent bactericidal activity with low resistance potential, yet their large-scale discovery from rapidly expanding genomic resources remains limited, as existing computational methods often rely on sequence homology and few reproducible, independently validated, and practically deployable prediction frameworks are currently available. We present LysinFusion, a reproducible deep learning framework that integrates heterogeneous sequence features via a hybrid CNN–Transformer architecture for accurate lysin identification. Trained on a curated, de-redundant corpus (PHROG + inphared) and validated on 148 experimentally confirmed UniProt proteins, LysinFusion achieves an accuracy of 0.81, an AUC of 0.89, and an MCC of 0.62, outperforming the state-of-the-art DeepMineLys by up to 52.3% across metrics while reducing false positives by 64% (12 vs. 33). Ablation studies confirm that both CNN and Transformer modules are individually essential and that serial processing outperforms parallel alternatives. Model interpretability via occlusion and LIME aligns with established lysin biology, with attention concentrating on the N-terminal catalytic domain and decision boundaries reflecting characteristic charge-composition thresholds documented in Gram-negative lysin C-terminal regions. The framework, including all source code, curated datasets, and an accompanying web server, is freely available at https://github.com/sinuo560/LysinFusion.

## 1. Introduction

The escalating antimicrobial resistance (AMR) crisis necessitates the urgent discovery of alternative antibacterial modalities beyond conventional small-molecule antibiotics^1^. Among the most promising candidates are phage lysins—peptidoglycan hydrolases that exhibit potent, species-specific bactericidal activity with a remarkably low propensity for resistance development^2–9^. Despite the exponential growth of phage genomic and metagenomic data, the large-scale mining of novel lysins remains hindered by inherent experimental bottlenecks, including the non-culturability of host microbes, holin-induced toxicity in expression systems, and the labor-intensive nature of functional screening^10–15^. Consequently, there is a critical demand for robust computational tools capable of accurately identifying lysins directly from amino acid sequences.

Early computational efforts employed pseudo-amino acid composition (PseAAC) coupled with support vector machines (SVM) or random forests^16,17^. While pioneering, these models were trained on limited datasets and, critically, many associated web servers—including Lypred—are no longer publicly accessible, severely hampering reproducibility and practical application^18–22^. Contemporary tools such as PhiBiScan and HMM-based strategies rely heavily on sequence homology searches against profile hidden Markov models (HMMs) derived from known lytic domains (e.g., PFAM, pVOGs)^23,24^. Although effective for conserved families, homology-driven approaches exhibit diminished sensitivity for highly divergent or novel lysins and impose substantial computational overhead when scanning terabase-scale metagenomic assemblies^24^.

Recent advances in deep learning, exemplified by DeepLysin and DeepMineLys, have demonstrated improved predictive power on unannotated open reading frames^25,26^. However, significant barriers to widespread adoption persist: DeepLysin lacks a publicly available implementation, limiting external validation; DeepMineLys, while performant, has not been evaluated on a fully independent, experimentally curated external benchmark, leaving questions regarding generalizability unaddressed^25,26^. Currently, the field lacks a fully reproducible, open-source, rigorously validated, and practically deployable deep learning framework for the binary classification of phage lysins. In many cases, previously published tools are either unavailable, difficult to reproduce, or lack accessible implementations and maintained web platforms, substantially limiting their long-term usability for large-scale lysin discovery^18–22,25,26^.

To bridge this gap, we introduce LysinFusion, a reproducible and high-fidelity framework for lysin identification. Leveraging a carefully curated, de-redundant training corpus derived from PHROG^27^, inphared^28^, and UniProt^29^, LysinFusion integrates four heterogeneous sequence descriptors (CKSAAP, CTDD, APAAC, CTDC) via a hybrid CNN-Transformer architecture. In rigorous independent validation against 148 experimentally confirmed lysins, LysinFusion achieves an accuracy of 0.81 and an AUC of 0.89, outperforming the state-of-the-art DeepMineLys by up to 52.3% across key metrics while reducing false positive identifications by 64%. Furthermore, we provide a user-friendly web server and fully containerized source code, establishing LysinFusion as a practical, reproducible, extensible, and transparent framework for high-throughput lysin candidate screening. By combining independent validation, biological interpretability, and public accessibility, LysinFusion addresses several longstanding limitations in existing lysin prediction tools and provides a deployable resource for large-scale genomic applications.

## 2. Materials and methods

### 2.1 Dataset

#### Training Set and validation set (PHROG + inphared)

The primary dataset used for model training and initial testing was constructed by combining protein sequences from two phage genome resources: PHROG^27^ and inphared^28^. PHROG provides high-quality, curated clusters of phage proteins but its latest update was in June 2022. To ensure that our model captured the most recent phage diversity, we supplemented PHROG with additional data downloaded from inphared, which is continuously updated.

For both sources, we first collected all protein sequences annotated with lysis-related functions as positive samples. To construct negatives with a consistent genetic background, we extracted the virus IDs of positive samples and then downsampled non-lysis proteins with the same virus IDs, excluding those annotated as “unknown” and retaining only those with clearly defined non-lysis functions.

We then performed several preprocessing steps to improve dataset quality. Sequences containing ambiguous amino acid letters were removed. Redundancy was reduced using CD-HIT. For positive lysin sequences, a high similarity threshold (-c 0.95 -n 5 - aS 0.8) was applied to retain the full evolutionary diversity of catalytic domains. For negative non-lysin sequences, a more stringent threshold (-c 0.4 -n 2 -aS 0.8) was applied to minimize redundancy among highly abundant phage structural and replication proteins, thereby enhancing the discriminatory challenge of the training data. After deduplication, positives were randomly split into training and validation sets at an 8:2 ratio, with negative samples subsequently balanced via random downsampling. In total, the final training set contained 18,865 sequences, and the validation set contained 4,717 sequences.

#### Independent Test Set (experimentally validated proteins from UniProt)

To more objectively assess the real-world performance of our model, we additionally built an independent test set consisting only of experimentally validated proteins. Many existing lysin prediction studies rely on random splits of collected data, which may include predicted or low-confidence annotations, potentially leading to overestimated performance. To avoid this bias, we curated a high-confidence benchmark set from the UniProtKB database^29^.

Specifically, we searched the UniProtKB database using the keyword phage and selected only the reviewed entries to ensure annotation reliability. Among the retrieved results, we retained entries with Protein existence levels of either “Protein level” (experimental evidence at the protein level) or “Transcript level” (experimental evidence at the transcript level). From this curated subset of experimentally supported phage-related proteins, we further filtered the entries using functional keywords such as endolysin, muramidase, amidase, CHAP, and peptidase to select lysin proteins as positive samples. All remaining entries within the same subset that did not contain any of these keywords were designated as non-lysin proteins and assigned as negative samples. After manual verification by two domain experts, the curated collection comprised 74 experimentally confirmed positive lysins and 1,286 high-confidence non-lysin proteins. To facilitate a balanced and conservative assessment of binary classification performance, a subset of 74 negative samples was randomly downsampled to constitute the final independent test set (1:1 ratio).

### 2.2 Sequence encoding schemes

We screened 29 amino acid sequence encoding schemes available in iLearn ^30^. For every encoder that could be executed on our datasets, we transformed each protein sequence into numerical features and evaluated a downstream classifier using the same protocol to ensure fair comparison.

For fast and consistent scoring, we used a LightGBM (LGBM) classifier, which trains quickly and outputs reliable probabilities^31^. On the validation set, we recorded accuracy (ACC), area under the ROC curve (AUC), Matthews correlation coefficient (MCC), F1-score (F1), sensitivity (SN), specificity (SP), runtime, and the feature dimensionality for each single encoder. We first ranked all single encoders by accuracy (ACC); if two encoders had the same ACC, we ranked them by area under the ROC curve (AUC). Encoders with very long runtime (> 1000s) were excluded from subsequent steps to speed up.

After this filtering, we performed a greedy combination search. Starting from the top-ranked single encoder, we iteratively added one remaining encoder at a time, concatenated feature blocks, retrained the same LGBM classifier, and kept the combination with the highest ACC (breaking ties by AUC). The search stopped once adding another encoder no longer improved ACC.

This process yielded a four-encoder feature set as our final scheme: CKSAAP + CTDD + APAAC + CTDC. In all subsequent experiments, we concatenate these four feature blocks for each sequence and use the combined vector for model training and evaluation.

### 2.3 Feature selection

We evaluated a broad set of feature selection (FS) strategies on top of the fixed four-encoder representation (CKSAAP + CTDD + APAAC + CTDC). For each FS method that could be executed, we applied it to the concatenated feature matrix from the training split, projected the same columns on the validation set, and trained the same downstream model to ensure comparability.

The scoring model was our CNN–Transformer architecture. Within each trial, features were standardized with StandardScaler, the network was trained on the training split with early stopping based on validation AUC, and then evaluated on the fixed test set. We recorded ACC, AUC, MCC, and F1 on the test set, and selected the FS method with the highest ACC as the final scheme.

The FS methods we tried covered common filters, wrappers, and embedded models, including but not limited to: low-variance / low non-zero-rate filtering (variance < 1e-6 or non-zero rate < 0.5%), correlation-based redundancy removal (absolute Pearson |r| ∈ {0.95, 0.97, 0.98, 0.99}), label correlation (point-biserial/Pearson) Top-K, univariate t-test / ANOVA-F Top-K, mutual information Top-K, chi-square Top-K (with MinMax scaling for scoring), LightGBM-based importance (Top-K and cumulative-gain thresholds), L1-regularized logistic regression with SelectFromModel, stability selection, and several chained pipelines (e.g., Var/NZ → MI → Corr, or Var/NZ → LGBM → Corr).

The final choice was the L1-regularized logistic regression selector with SelectFromModel (penalty=L1, solver=liblinear, C=0.1), preceded by a fast prefilter that removed features with variance < 10^−6^ or non-zero rate < 0.5%. After selection, we zero-padded the feature dimension to a multiple of 400 to match the CNN input reshaping used by our model. In all subsequent experiments, we applied this FS chain to the four-encoder features and used the selected columns for model training and evaluation.

### 2.4 General framework of LysinFusion

Figure 1 overviews the full architecture of the LysinFusion framework. Starting from labeled protein sequences, four complementary sequence descriptors are computed: CKSAAP (gap = 5), CTDD, APAAC (w = 0.05, lambda adaptive to sequence length), and CTDC. These are concatenated into a single feature vector per sequence. To ensure compatibility between training and test sets, encoder outputs are aligned in length via padding or truncation, and all numerical anomalies are sanitized to zero.

**Figure 1:**
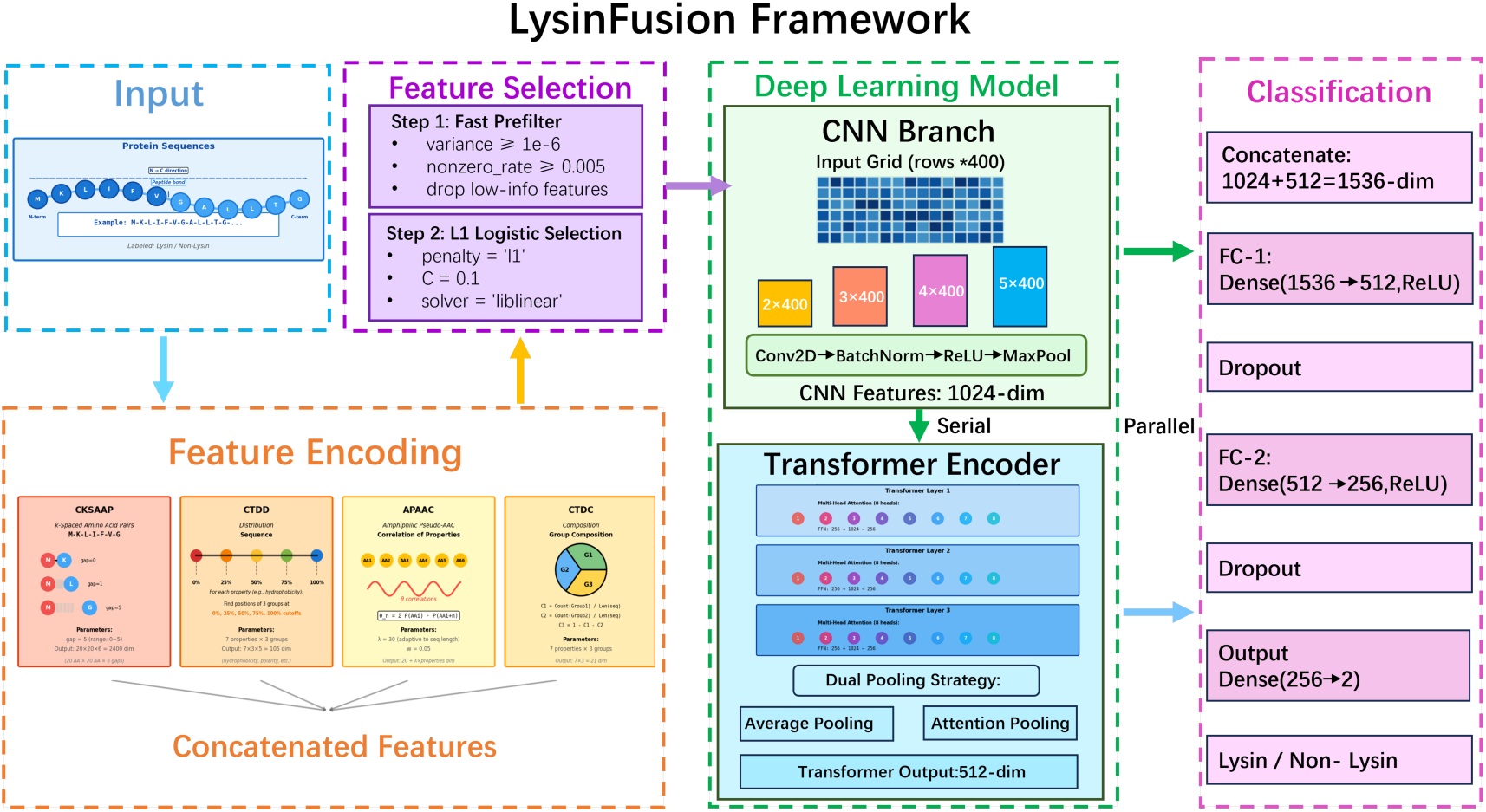
The pipeline integrates four sequence encoders, a two-step feature selection process, and a hybrid architecture that combines sequential CNN–Transformer modeling with parallel feature fusion for final classification.

We then apply a two-step feature selection procedure. A statistical pre-filter removes features with low variance or very low non-zero rates. Second, L1-regularized logistic regression is used with SelectFromModel (C=0.1, penalty=l1, solver=liblinear) to identify discriminative features. If no features are retained, the system falls back to selecting features with the largest coefficient magnitudes. The final feature vector is zero-padded to a minimum dimension of 2000 and adjusted to a multiple of 400, then reshaped into a 2D grid format (grid rows × 400, where grid rows ≥ 5) as input to the deep model.

The classifier combines both serial and parallel architectural elements. The reshaped input grid is first passed through a 2D convolutional encoder (TextCNN) with kernel heights of 2, 3, 4, and 5, each with 256 channels. After batch normalization, ReLU activation, max-pooling, and dropout, a dense layer reduces the features to a 1024-dimensional vector with tanh activation.

This CNN output is then transformed into a sequence representation for the Transformer encoder. Specifically, it is projected to 400 dimensions, combined with positional embeddings to form a sequence of length 60, then projected again to 256 dimensions with scaling and additional positional encodings. The resulting sequence is processed through a 3-layer Transformer encoder with 8 attention heads per layer, along with residual connections and layer normalization.

To capture both global and local context, we extract two representations from the Trans-former output: average-pooled and soft attention-pooled embeddings. These are concatenated with the original CNN features in a parallel fusion step. The combined vector is then passed through a two-layer fully connected head (FC-512 → dropout → FC-256 → dropout → 2-way logits) for final classification.

Training is conducted with fixed training and test sets. Features are standardized using StandardScaler, with 10 percent of the training set held out for validation. The model is optimized using Adam with exponential learning rate decay (rate 0.95 per 1000 steps), L2 regularization, and global gradient clipping at 5.0. Early stopping is triggered if validation AUC fails to improve for 5 consecutive evaluations (evaluated every 5 epochs). At inference, predictions are made by thresholding the positive-class softmax probability at 0.5. Evaluation metrics include accuracy, AUC, MCC, and F1 score.

### 2.5 Occlusion-based interpretability analysis

To evaluate the contribution of different regions within protein sequences to the prediction performance of our classification model, we adopted an occlusion-based interpretability analysis using a sliding window strategy. This method involves systematically masking specific segments of the amino acid sequence and comparing model performance before and after occlusion, thereby identifying the most influential positions. The analysis was adapted from the masking-based interpretability strategy used in our previous study^32^, where a sliding mask was applied to biological sequences to evaluate the contribution of different sequence regions to model prediction. In the present work, we modified this strategy for phage lysin protein sequences by grouping proteins according to sequence length, masking amino acid segments with “X”, applying multiple window sizes, and evaluating the resulting performance degradation using the LysinFusion feature-encoding and CNN–Transformer framework. These modifications were introduced to better accommodate the large variation in protein sequence lengths and the more complex structural organization of phage lysins compared with the RNA sequences analyzed in our previous work.

#### Data Preprocessing

We first merged the de-redundant positive and negative protein sequences collected from both the PHROG database and the inphared resource, and then calculated the sequence lengths for each sample. To handle large variations in sequence length, we binned sequences into fixed-length intervals. The distribution showed that most sequences fell within the 0–400 amino acid range, including approximately 176,600 sequences in the 0–200 interval (48.4%) and 115,800 sequences in the 200–400 interval (31.7%), accounting for about 80% of all samples. Sequences outside this range were far fewer. Based on this distribution, we selected sequences within the 0–200 and 200–400 amino acid length ranges as representative subsets for subsequent analysis. To ensure consistency across occlusion and control experiments, each subset was randomly shuffled once, and a unique ID was assigned to every sequence. This order and ID mapping were preserved throughout all experiments to strictly control for variables other than the occluded positions.

#### Occlusion Experiment Design and Model Evaluation

We applied a sliding window occlusion strategy to simulate the effect of masking specific sequence regions on model performance. Multiple window lengths were defined for each sequence length group. For both the 0–200 and 200–400 groups, window lengths k ranged from 10 to 100 (step size = 10). The same range of window sizes was applied to ensure comparability of occlusion effects across different sequence length groups. At each k value, the window slid from the beginning to the end of each sequence, shifting one residue at a time. Sequences shorter than the current occlusion window were padded with “X” at the end to enable occlusion. Within each window, the original amino acids were replaced with “X”, which represents an unknown residue and effectively masks sequence information.

After occlusion, sequences were split into training and test sets using a fixed 8:2 ratio. Each sequence was encoded using the four selected descriptors (CKSAAP, CTDD, APAAC, and CTDC), and the LysinFusion model was trained for classification. For each experiment, we recorded the window length, starting position, and four performance metrics: Accuracy, Matthews Correlation Coefficient (MCC), Area Under the ROC Curve (AUC), and F1 score.

#### Control Experiment

To provide a baseline for comparison, we performed control experiments without any occlusion. Each subset was split into training and test sets using the same sample order and 8:2 ratio as in the occlusion experiments. The same four encodings and the LysinFusion model were used. Model performance on the test set was recorded using the same four metrics: Accuracy, MCC, AUC, and F1 score. These control results served as the reference for measuring performance degradation due to occlusion.

#### Performance Delta Calculation

For each occlusion experiment, we calculated the performance degradation (delta) for the four metrics by subtracting the occluded performance from the control performance. Delta values reflect the degree to which masking a specific region reduces model accuracy and thus indicate the importance of that region. We then mapped the occlusion window to individual amino acid positions and assigned the delta value to all affected positions. For each position, we aggregated all associated delta values and computed the average delta per metric, thereby producing a position-wise impact profile for all four performance indicators. This position-wise impact profile was subsequently visualized to identify the most influential regions contributing to model predictions.

### 2.6 Local interpretable model-agnostic explanations (LIME)

Deep learning models, while powerful, are often regarded as “black boxes” because their internal decision process is difficult to interpret. To address this issue, we adopted Local Interpretable Model-agnostic Explanations (LIME)^33^, which builds a simple local model around each prediction to explain how input features contribute to that specific output.

To systematically interpret our model’s decision process, we first encoded each protein sequence in the training set using four descriptors (CKSAAP, CTDD, APAAC, and CTDC) to generate feature matrices. These encoded blocks were concatenated and further refined through a two-step feature selection process to remove redundant or uninformative variables. The selected features were then used as input to train our framework model. After training, the model was applied to the independent test set of 148 sequences to obtain prediction probabilities for both classes. Each test sequence was then categorized into one of four groups—true positive (TP), true negative (TN), false positive (FP), or false negative (FN)—based on the comparison between predicted and true labels. This grouping allowed us to separately analyze correctly classified lysins and non-lysins.

For each correctly predicted sample (TP and TN), we used LIME to generate feature-level explanations. Concretely, LIME creates many small perturbations of the target sample to form a local neighborhood, fits a simple linear surrogate to approximate our model’s behavior in that neighborhood, and uses the surrogate coefficients as feature contributions. Positive coefficients indicate features that support the predicted class, whereas negative coefficients indicate features that oppose it. By repeating this process across all correctly predicted samples, we obtained a collection of local feature importance scores. These scores were then combined to reveal overall patterns of feature relevance across the test set, linking local explanations to the model’s global decision behavior.

To illustrate this process and provide an example of the generated explanations, we visualized one representative prediction (Fig. 2). The left panel shows the prediction probabilities for the two classes, where the model assigns a probability of 0.98 to the negative class and 0.02 to the positive class. The central panel highlights the most influential features contributing to this decision, with blue bars supporting the negative prediction and orange bars favoring the positive prediction. The magnitude of each bar reflects the weight of the corresponding feature in the local surrogate model. Finally, the right panel lists the values of these high-impact features for the specific sample. This visualization reveals not only the predicted label but also the feature-level reasoning behind it, thereby improving the interpretability and trustworthiness of our model’s decisions.

**Figure 2:**
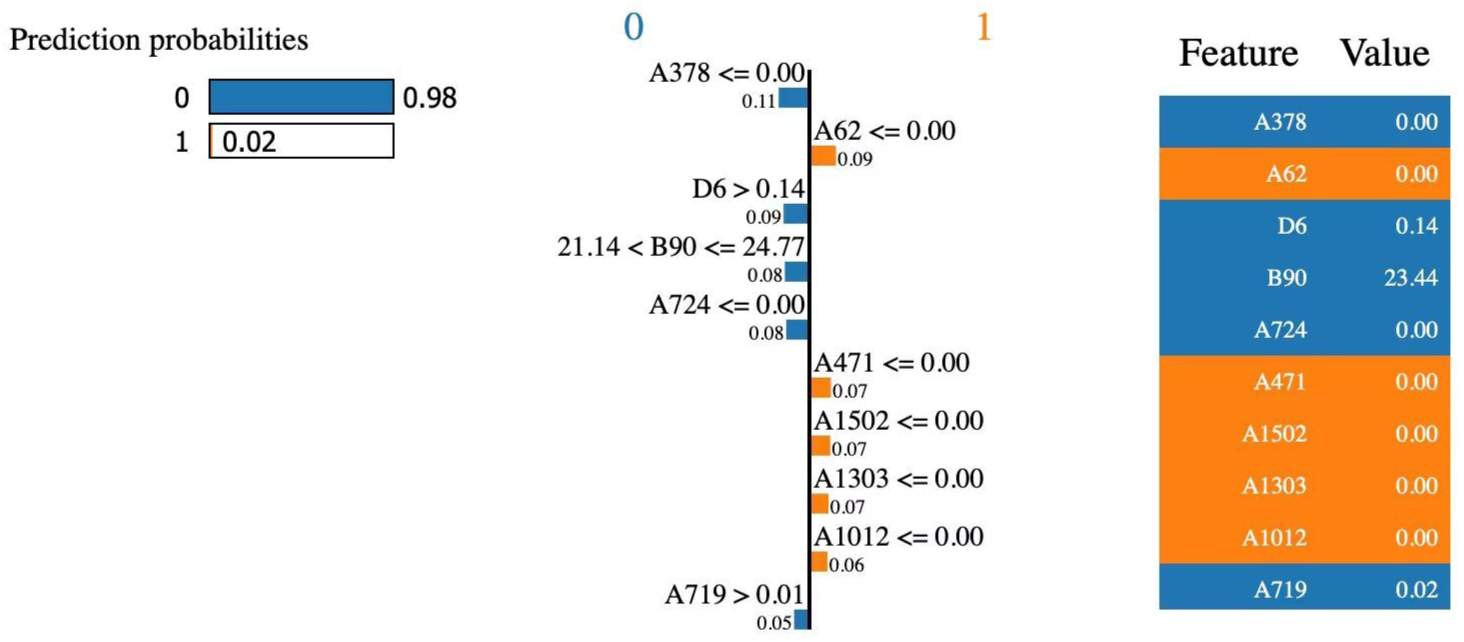
LIME-based explanation of a representative prediction. Left: prediction probabilities for the two classes. Middle: most influential features and their contribution directions (blue: negative class, orange: positive class). Right: actual values of these features in the analyzed sample.

### 2.7 Performance evaluation metrics

To rigorously assess the classification performance of different sequence encoding methods, we employed six widely recognized evaluation metrics: Accuracy (ACC), Sensitivity (SN), Specificity (SP), Matthews Correlation Coefficient (MCC), F1-score (F1), and the Area Under the Receiver Operating Characteristic Curve (AUC). These metrics are essential for providing a holistic evaluation of model performance, including its correctness, robustness on imbalanced datasets, and general classification capability.

The mathematical definitions of these metrics are expressed as follows:

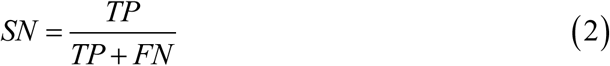

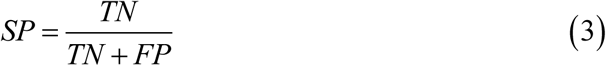

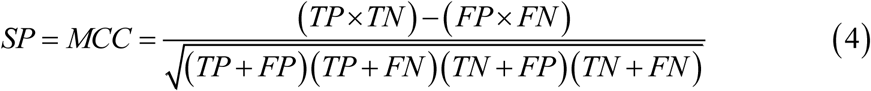

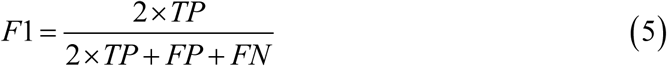

In these formulas, *TP*, *TN*, *FP*, and *FN* denote true positives, true negatives, false positives, and false negatives, respectively. *TP* represents the count of positive samples correctly identified by the model, while *TN* corresponds to negative samples correctly predicted. Conversely, *FP* and *FN* reflect incorrect predictions where negative samples are misclassified as positive and positive samples are misclassified as negative, respectively.

The Area Under the Receiver Operating Characteristic Curve (AUC) is an essential threshold-independent evaluation metric. It reflects the probability that a randomly chosen positive sample is ranked higher than a randomly selected negative sample by the classifier. Mathematically, AUC can be calculated using the following formula^34^:

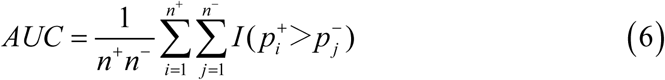

where *n*^+^ and *n*^−^ denote the number of positive and negative samples, respectively. 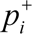 and 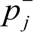 are the prediction scores assigned to the *i*-th positive and *j*-th negative samples. The indicator function *I*(·) returns 1 if 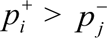, and 0 otherwise. This formulation effectively quantifies the ranking capability of the classifier over the entire dataset without depending on a specific decision threshold

By utilizing these six metrics, we conducted a comprehensive evaluation of model performance, ensuring balanced consideration of overall accuracy, sensitivity towards detecting positive instances, specificity for identifying negative samples, and threshold-free classification ability through *AUC*.

## 3. Results and discussion

### 3.1 Evaluation of various sequence encoding methods

Protein sequences can be represented by many encoding schemes, and choosing an effective encoder is essential. We evaluated 29 commonly used methods available in iLearn: CKSAAP, TPC, DDE, DPC, APAAC, PAAC, QSOrder, CTDD, EAAC, CKSAAGP, AAC, BLOSUM62, ZSCALE, EGAAC, CTDDClass, Binary, GTPC, CTDC, NMBroto, CTriad, KSCTriad, CTDT, Geary, Moran, CTDTClass, GDPC, SOCNumber, CTDC-Class, and GAAC. Each encoder was applied once to transform sequences, and the same downstream LightGBM classifier and the validation set were used for scoring (ACC, AUC, MCC, F1, SN, SP); ranking was by ACC. Figure 3 (ACC) and Figure 4 (AUC) visualize the single-encoder comparison. CKSAAP was the most effective encoder, reaching an ACC of **0.9350** and an AUC of **0.9797**. TPC and DDE also performed strongly (both ACC > 0.91; TPC AUC ≈ 0.972, DDE AUC ≈ 0.969).

**Figure 3:**
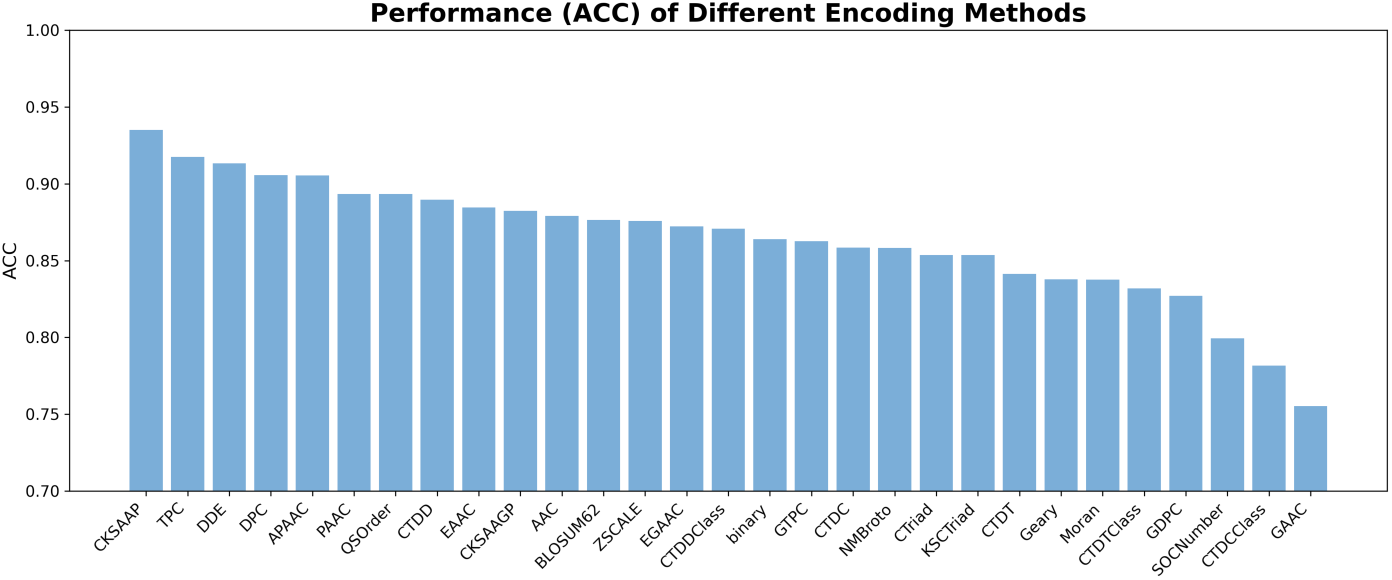
Test accuracy (ACC) of the 29 single encoders.

**Figure 4:**
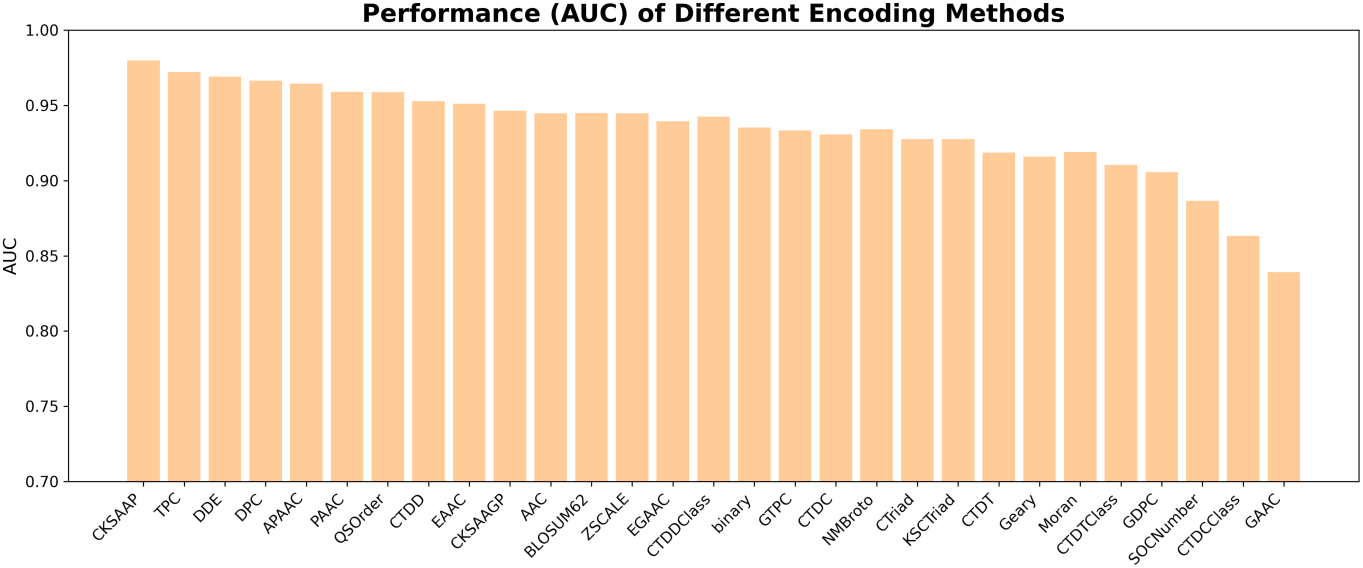
Test AUC of the 29 single encoders.

However, TPC required a significantly longer runtime (approximately 4.4 hours, see Figure 5) compared to other methods (median runtime: 68.6 seconds) and was therefore excluded from later combination steps to keep the pipeline efficient. Guided by this ranking, we then conducted a greedy combination search: starting from the top single encoder CKSAAP, we added one remaining encoder at a time, concatenated the feature blocks, retrained the same classifier, and kept the best ACC in each round (if ACC tied, we compared AUC). The trajectory is shown in Figure 6. With CKSAAP as the baseline (Round 0), performance increased monotonically in the first three rounds: Round 1 (CKSAAP+CTDD) achieved ACC=0.9410 and AUC=0.9822; Round 2 (CKSAAP+CTDD+APAAC) further improved to ACC=0.9444 and AUC=0.9837; Round 3(CKSAAP+CTDD+APAAC+CTDC) reached the best ACC=0.9464 with AUC=0.9839. Adding a fifth block (Round 4, +DPC) did not improve ACC (still 0.9464), indicating that the marginal benefit saturates once four complementary encoders are combined. Considering the diminishing returns, the increased dimensionality, and the runtime budget. Consequently, the integration of CKSAAP, CTDD, APAAC, and CTDC yielded the optimal balance between predictive power and feature dimensionality, with no further gains observed upon the inclusion of a fifth descriptor (ACC remained at 0.9464).

**Figure 5:**
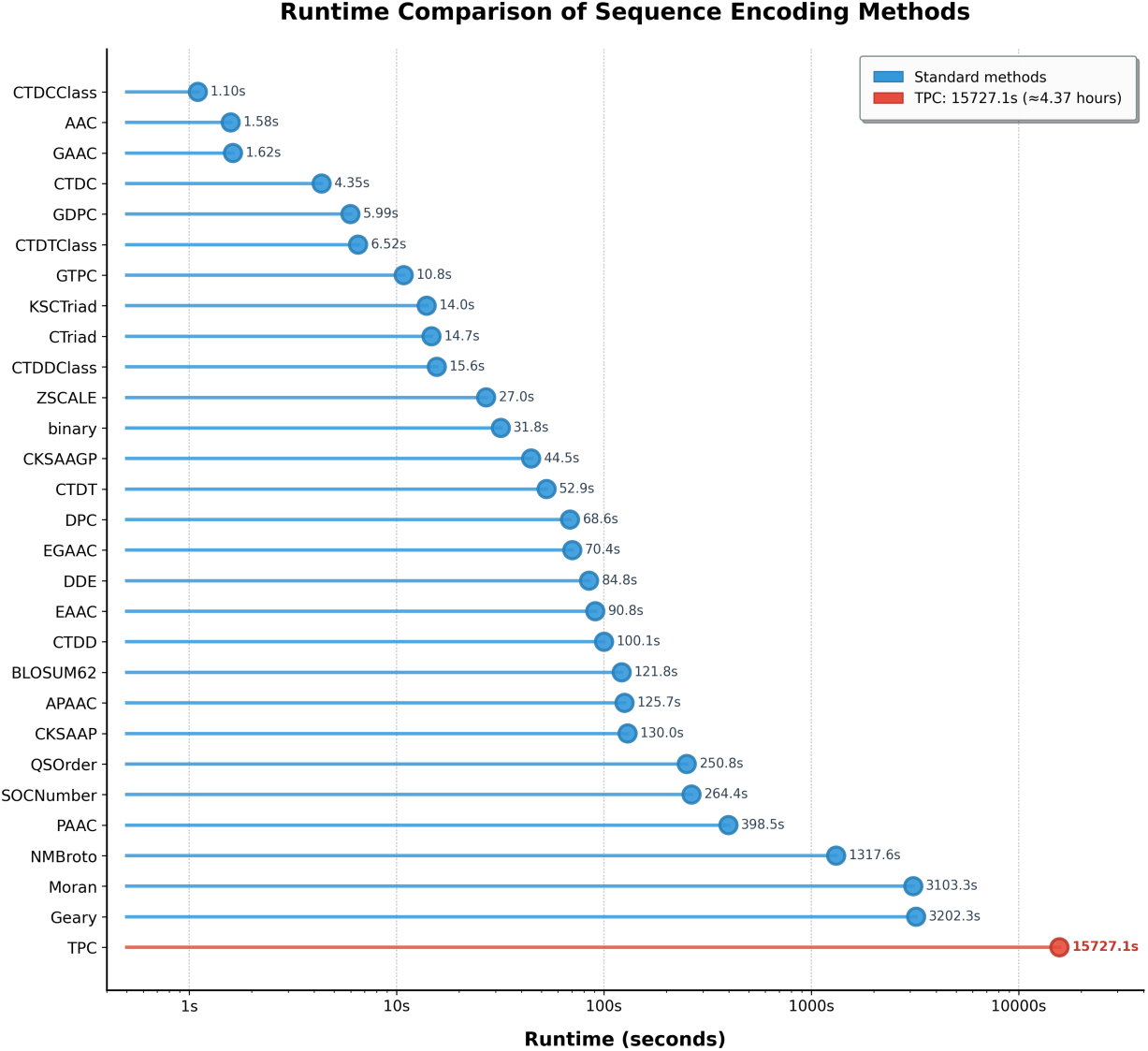
Runtime comparison of sequence encoding methods. Methods are sorted by runtime (slowest to fastest). TPC is an extreme outlier, requiring approximately 4.4 hours, while most methods complete within minutes. The fastest method (CTDCClass) takes only 1.1 seconds.

**Figure 6:**
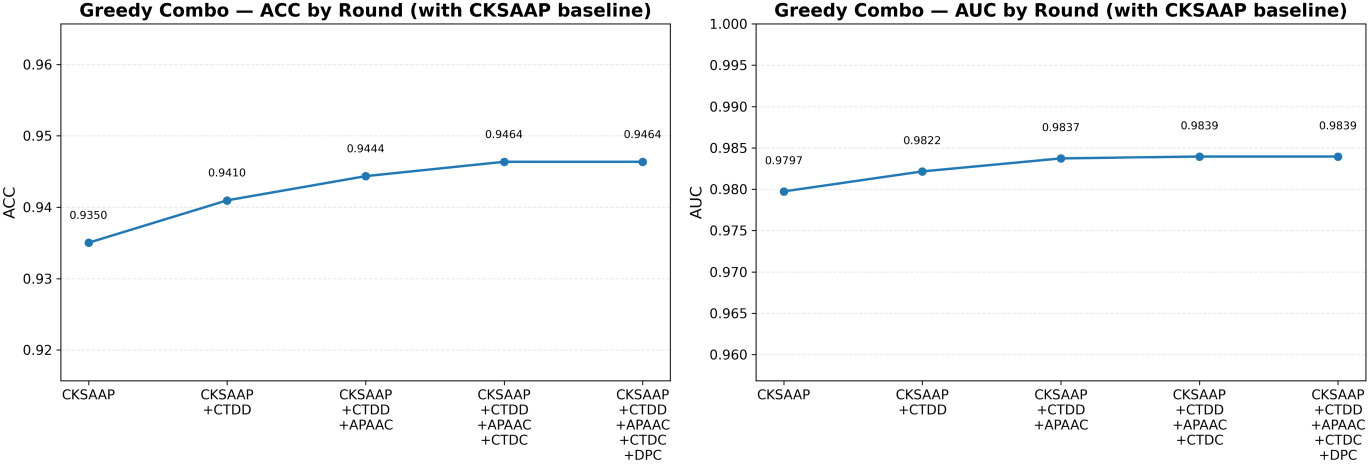
Greedy combination search starting from CKSAAP (Round 0).

### 3.2 Evaluation of various feature selection methods

We benchmarked a range of feature selection strategies applied to the concatenated four-encoder representation, evaluating both predictive performance and dimensionality reduction efficiency (Table 1).

**Table 1:** Comparison of feature selection methods on the four-encoder feature set. All methods use the same train/validation split and classifier. Best results are highlighted in bold.

| Feature Selection Method | Parameters | Selected Dim | ACC | AUC | MCC | F1 |
| --- | --- | --- | --- | --- | --- | --- |
| Var/NZ + Corr | var=1e-6, nz=0.5%, r=0.98 | 2686 | 0.9438 | 0.9838 | 0.8876 | 0.9437 |
| Var/NZ-only | var=1e-6, nz=0.5% | 2710 | 0.9483 | 0.9847 | 0.8965 | 0.9482 |
| Corr-only (inter-feature redundancy removal) | r=0.95 | 2676 | 0.9413 | 0.9822 | 0.8834 | 0.9399 |
| Corr-only (inter-feature redundancy removal) | r=0.97 | 2683 | 0.9471 | 0.9837 | 0.8944 | 0.9465 |
| Corr-only (inter-feature redundancy removal) | r=0.98 | 2686 | 0.942 | 0.9822 | 0.884 | 0.9419 |
| Corr-only (inter-feature redundancy removal) | r=0.99 | 2687 | 0.9492 | 0.9847 | 0.8991 | 0.9482 |
| Var/NZ + Corr | var=1e-6, nz=0.5%, r=0.95 | 2676 | 0.9424 | 0.9847 | 0.885 | 0.943 |
| Var/NZ + Corr | var=1e-6, nz=0.5%, r=0.97 | 2683 | 0.9421 | 0.9826 | 0.8856 | 0.9405 |
| Var/NZ + Corr | var=1e-6, nz=0.5%, r=0.99 | 2687 | 0.943 | 0.985 | 0.886 | 0.9432 |
| Label Correlation (Pearson) Top-K | K=512 (point-biserial correlation for $y \in \{0,1\}$) | 2000 | 0.9352 | 0.9776 | 0.8717 | 0.9335 |
| Label Correlation (Pearson) Top-K | K=1024 | 2000 | 0.9403 | 0.9814 | 0.8807 | 0.9406 |
| Label Correlation (Pearson) Top-K | K=2048 | 2048 | 0.9469 | 0.9849 | 0.8938 | 0.9466 |
| Label Correlation (Pearson) Top-K | K=4096 | 2710 | 0.949 | 0.9861 | 0.8981 | 0.9494 |
| T-test/ANOVA-F Top-K | K=512 | 2000 | 0.9325 | 0.9789 | 0.8651 | 0.9331 |
| T-test/ANOVA-F Top-K | K=1024 | 2000 | 0.9406 | 0.9828 | 0.8813 | 0.9407 |
| T-test/ANOVA-F Top-K | K=2048 | 2048 | 0.9487 | 0.9866 | 0.8979 | 0.9478 |
| T-test/ANOVA-F Top-K | K=4096 | 2710 | 0.9476 | 0.9861 | 0.8953 | 0.9478 |
| Mutual Information (MI) Top-K | K=512 | 2000 | 0.9372 | 0.9804 | 0.8745 | 0.937 |
| Mutual Information (MI) Top-K | K=1024 | 2000 | 0.9353 | 0.9802 | 0.8707 | 0.9356 |
| Mutual Information (MI) Top-K | K=2048 | 2048 | 0.9365 | 0.9833 | 0.8734 | 0.9374 |
| Mutual Information (MI) Top-K | K=4096 | 2710 | 0.9458 | 0.986 | 0.8917 | 0.9462 |
| Chi-square ($\chi^2$) Top-K | K=1024 (MinMax[0,1] normalization during scoring) | 2000 | 0.9335 | 0.9793 | 0.8673 | 0.9327 |
| Chi-square ($\chi^2$) Top-K | K=2048 (MinMax[0,1] normalization) | 2048 | 0.9447 | 0.9845 | 0.8894 | 0.9442 |
| LightGBM Feature Importance Top-K | K=1024, n estimators≈200, lr≈0.05 | 2000 | 0.9465 | 0.9843 | 0.8929 | 0.9464 |
| LightGBM Cumulative Importance | Cumulative gain≥95%, n estimators≈200 | 2000 | 0.9379 | 0.9846 | 0.8763 | 0.939 |
| **L1 Logistic (SelectFromModel)** | **C=0.1, penalty=L1, solver=liblinear** | **2243** | **0.9506** | 0.9851 | **0.9012** | **0.9506** |
| L1 Logistic (SelectFromModel) | C=1.0, penalty=L1, solver=liblinear | 2541 | 0.9497 | **0.9867** | 0.8995 | 0.9491 |
| Combined: Var/NZ → MI → Corr | var=1e-6, nz=0.5%, K=2048, r=0.98 | 2024 | 0.9417 | 0.9836 | 0.8834 | 0.9416 |
| Combined: Var/NZ → LGBM → Corr | var=1e-6, nz=0.5%, K=2048, r=0.98 | 2026 | 0.9497 | 0.9865 | 0.8995 | 0.9491 |
| Block Fair Sampling → Corr | top-p%=20% per encoding block, r=0.98 | 2000 | 0.9283 | 0.9754 | 0.8568 | 0.9278 |
| Stability Selection (MI-based) | seeds=5, K=2048 each run, keep≥60%, followed by r=0.98 | 2028 | 0.9392 | 0.983 | 0.8785 | 0.9398 |

Across all methods, the L1-regularized logistic regression selector (SelectFromModel) was the best overall. With C = 0.1 it achieved the highest test accuracy, ACC = 0.9506, together with MCC = 0.9012 and F1 = 0.9506, while reducing the representation to 2,243 dimensions. The C = 1.0 variant reached the top AUC (AUC = 0.9867) with very similar accuracy (ACC = 0.9497; 2,541 dims), confirming the robustness of the L1 embedded selector.

Several simple filters were competitive but slightly behind: correlation de-redundancy with a strict threshold (r = 0.99) reached ACC = 0.9492 (2,687 dims); label-correlation Top-K peaked at K = 4096 with ACC = 0.9490 and AUC = 0.9861 (2,710 dims); ANOVA-F Top-K at K = 2048 obtained ACC = 0.9487 and AUC = 0.9866 (2,048 dims). A fast variance/non-zero filter alone already gave ACC = 0.9483 (2,710 dims). LightGBM importance Top-K yielded ACC = 0.9465 (2,000 dims). Methods based on mutual information and χ^2^ were moderate at best (best MI: ACC = 0.9458; best χ^2^: ACC = 0.9447), and the block-wise fair-sampling heuristic was clearly worse (ACC = 0.9283).

Ultimately, we chose **L1-logistic** (*C* = 0.1) **with a lightweight Var/NZ prefilter** as the final FS module: it delivers the best accuracy and the strongest MCC/F1 while trimming the feature space to ∼ 2.2k dimensions, striking a favorable balance between performance and compactness for all subsequent experiments.

### 3.3 Ablation study

To quantify the contribution of each main component in our pipeline, we conducted an ablation study on an independent test set of 148 sequences. The **complete model** uses four concatenated protein encodings (CKSAAP, CTDD, APAAC, CTDC), L1-regularized logistic regression for feature selection, and a hybrid architecture where: (1) CNN extracts local features, (2) these CNN features are mapped to a sequence and fed into a Trans-former, and (3) both CNN features and Transformer outputs are concatenated for final classification.

We systematically removed one component at a time to create five ablated variants: (1) **No Feature Selection**, which skips feature selection and uses all 2682 concatenated features directly; (2) **No CNN**, which removes the CNN module and lets the Transformer directly process the input features; (3) **No Transformer**, which removes the Transformer module and classifies using only CNN features; (4) **No Skip Connection**, which keeps the CNN→Transformer serial flow, but the final classifier uses only Transformer output without concatenating CNN features; and (5) **Parallel Only**, where CNN and Transformer independently process the input in parallel (rather than serially), and their outputs are concatenated for classification.

As illustrated in the radar chart (Fig. 7), which simultaneously compares all configurations across four key metrics, the complete model achieves ACC 0.8108, AUC 0.8921, MCC 0.6225, and F1 0.8056. Removing feature selection causes a minor drop (ACC −0.7%, MCC −1.4 points), confirming that feature selection improves robustness. Removing the CNN module (No CNN) leads to the most severe degradation (ACC −4.7%, MCC −9.5 points), demonstrating that CNN-extracted local motifs are critical. Removing the Transformer (No Transformer) also substantially hurts performance (ACC −2.7%, MCC−5.5 points), showing that global contextual modeling is essential.

**Figure 7:**
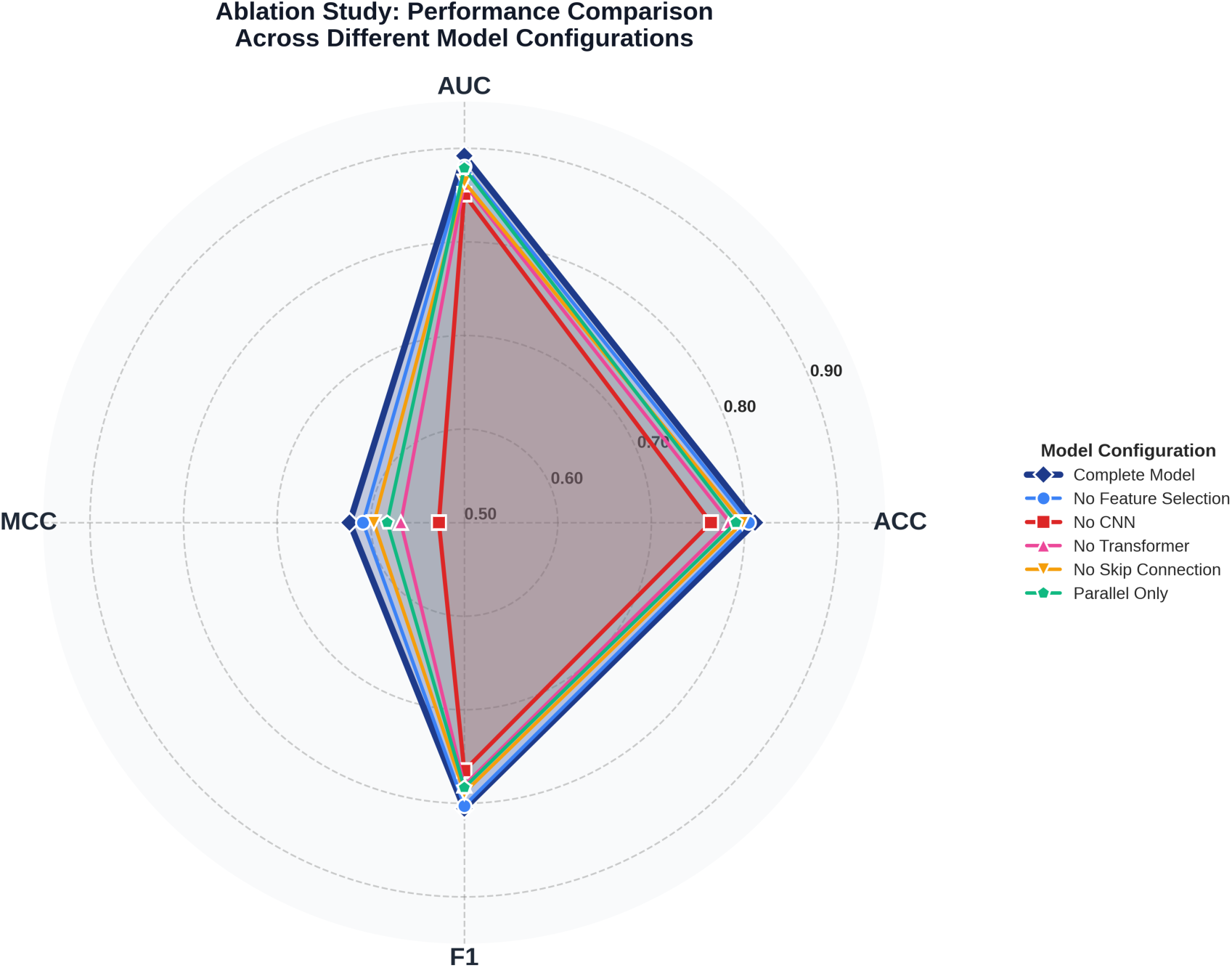
Ablation study results on the independent test set (148 sequences). The radar chart visualizes the multi-dimensional performance comparison across four evaluation met-rics (ACC, AUC, MCC, F1) for the complete model and five ablated variants. The com-plete model (outermost polygon) demonstrates superior performance across all metrics.

The No Skip Connection variant drops moderately (ACC −1.4%, MCC −2.6 points), indicating that directly passing CNN features to the classifier—bypassing the Trans-former—retains useful information. Finally, Parallel Only underperforms the complete model (ACC −2.0%, MCC −4.0 points), revealing that the serial architecture (CNN–Transformer) is superior to parallel processing. The serial design allows the Transformer to build upon CNN-extracted features rather than processing raw inputs independently. Overall, all five components contribute meaningfully to the model’s performance.

### 3.4 Comparative evaluation with existing methods

To comprehensively evaluate the predictive capability of our framework, we compared it with *DeepMineLys*^26^, which is the only publicly available and reproducible deep learning-based method for lysin prediction. Other existing approaches, such as PhiBiScan or the recently published curated HMM collections, rely on profile similarity searches rather than classification models, and their results are strongly dependent on user-defined model sets and score thresholds. These characteristics make direct comparison difficult. Therefore, we only compared our model with DeepMineLys. This limitation itself reflects the current lack of reproducible, accessible, and standardized benchmarking frameworks in the lysin prediction field.

For a fair and unbiased evaluation, both methods were tested on the same independent test set, which contains 148 experimentally validated protein sequences (74 lysins and 74 non-lysins). We followed the official DeepMineLys inference procedure to obtain its predictions and then applied our proposed LysinFusion model to the same dataset. Model performance was assessed using Accuracy (ACC), Sensitivity (SN), Specificity (SP), Matthews Correlation Coefficient (MCC), Area Under the ROC Curve (AUC), and F1-score (F1).

As illustrated in the dumbbell chart (Fig. 8), which highlights the performance gap between the two models across six evaluation metrics, LysinFusion demonstrates clear and consistent improvements over DeepMineLys across most key metrics. Our model achieves an ACC of 0.8108, representing a 16.5% improvement, and an AUC of 0.8921, reflecting a 19.5% gain in overall discriminative ability. The MCC of 0.6225 is more than 50% higher than that of DeepMineLys, highlighting the superior balance of positive and negative pre-dictions. In terms of classification behavior, LysinFusion produced only 12 false positives compared with 33 from DeepMineLys, indicating that our model greatly reduces spurious predictions. Although LysinFusion shows slightly lower sensitivity (78.38% vs. 83.78%), this minor reduction in recall is compensated by a much higher specificity (83.78% vs. 55.41%), resulting in a more reliable predictor that effectively limits false alarms. Consequently, our model also achieves a higher F1-score (0.8056 vs. 0.7337), confirming its better overall balance between precision and recall.

**Figure 8:**
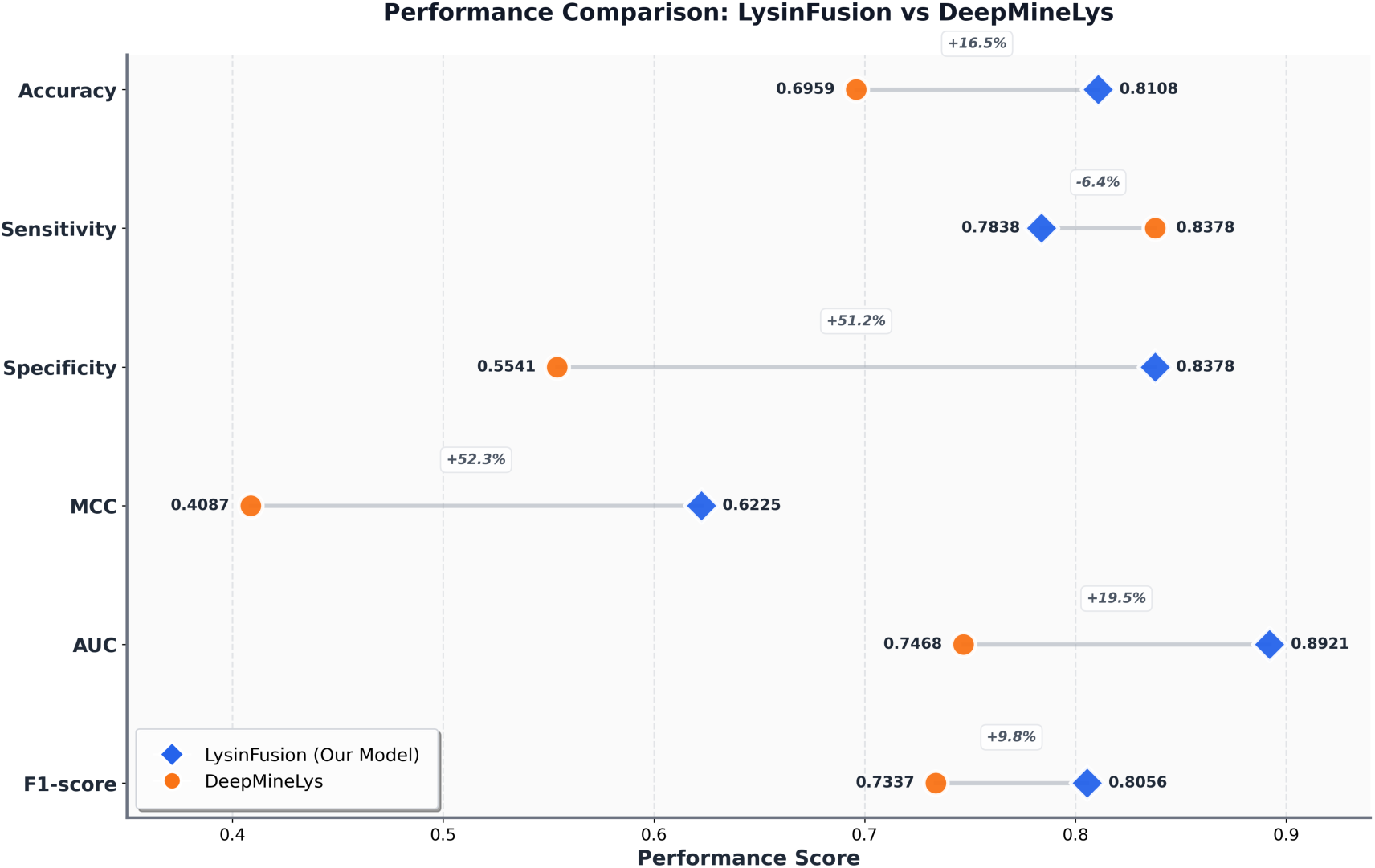
Performance comparison between LysinFusion and DeepMineLys on indepen-dent test set. The dumbbell chart visualizes the performance gap across six evaluation metrics, with connecting lines indicating the magnitude of difference. LysinFusion (blue diamonds) achieves consistently higher accuracy, specificity, MCC, AUC, and F1-score, while DeepMineLys (orange circles) shows slightly higher sensitivity.

From a practical perspective, the high specificity and low false-positive rate of LysinFusion are particularly valuable for real-world applications, where experimental validation is costly and time-consuming. In realistic protein screening scenarios, lysins represent only a very small fraction of the proteome, so keeping the number of incorrect predictions low is crucial to avoid overwhelming the true hits. By reducing the number of false candidates by nearly two-thirds, LysinFusion can significantly lower the experimental workload and improve the efficiency of lysin discovery pipelines. Overall, these results demonstrate that LysinFusion is a more accurate, robust, and cost-effective solution compared with the current state-of-the-art.

### 3.5 Interpretability analysis

To interpret how LysinFusion makes its predictions, we analyzed the model from two complementary perspectives. The first focuses on sequence regions, identifying where along the protein the model derives its main predictive information using an occlusion-based approach. The second examines the contribution of physicochemical and compositional features through LIME. Together, these analyses provide an overview of both positional and feature-level factors influencing the model’s classification decisions.

#### 3.5.1 Occlusion-based results

We next examined the results of the occlusion-based interpretability analysis to identify which sequence regions contribute most strongly to the predictive performance of the LysinFusion model. By sliding a masking window across protein sequences and measuring the resulting performance degradation, we obtained position-wise impact profiles for different sequence length ranges. The aggregated results are summarized below.

For sequences with lengths in the 0–200 range (Figure 9), performance deltas were highest in the early positions. In particular, F1 score and MCC showed a sharp peak between positions 10 and 25, indicating that masking residues in this region led to the most severe drops in predictive power. AUC also displayed a moderate elevation in this region, while accuracy remained relatively stable with only minor fluctuations. Beyond approximately position 100, the deltas of all metrics declined toward zero, suggesting that later positions in shorter sequences contributed less to the model’s predictions.

**Figure 9:**
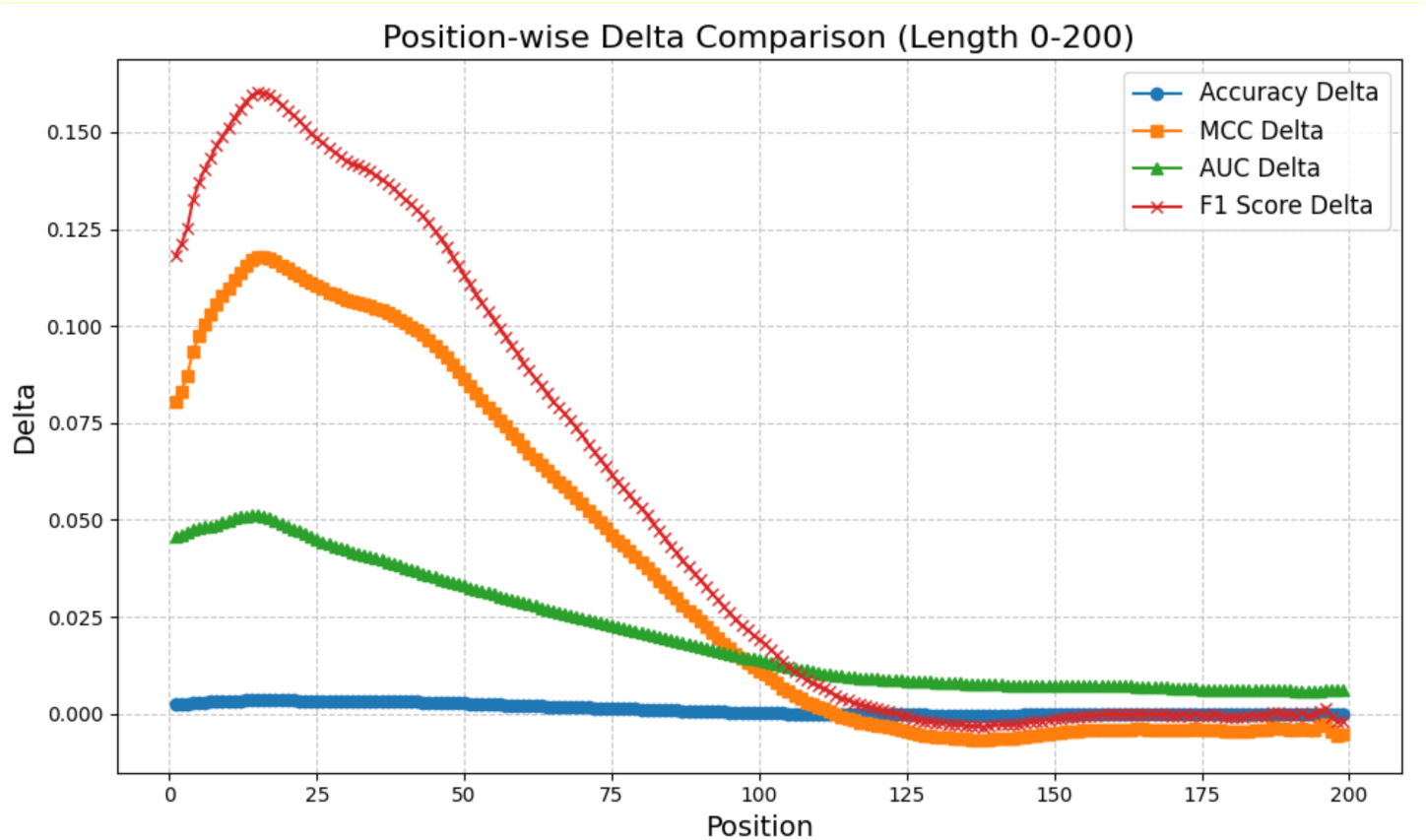
Position-wise delta comparison for sequences of length 0–200. F1 score and MCC show the highest sensitivity to masking in early positions, while accuracy remains nearly flat.

For the 200–400 range (Figure 10), the trends were similar but more pronounced. The early positions again exhibited the strongest performance degradation, with F1 score deltas exceeding 0.35 and MCC deltas close to 0.30. AUC started around 0.16 and decreased steadily across positions, while accuracy deltas remained nearly flat at close to zero. Importantly, the impact of occlusion diminished gradually across positions but stayed positive throughout most of the sequence, indicating that longer proteins contain more distributed yet still front-loaded informative regions.

**Figure 10:**
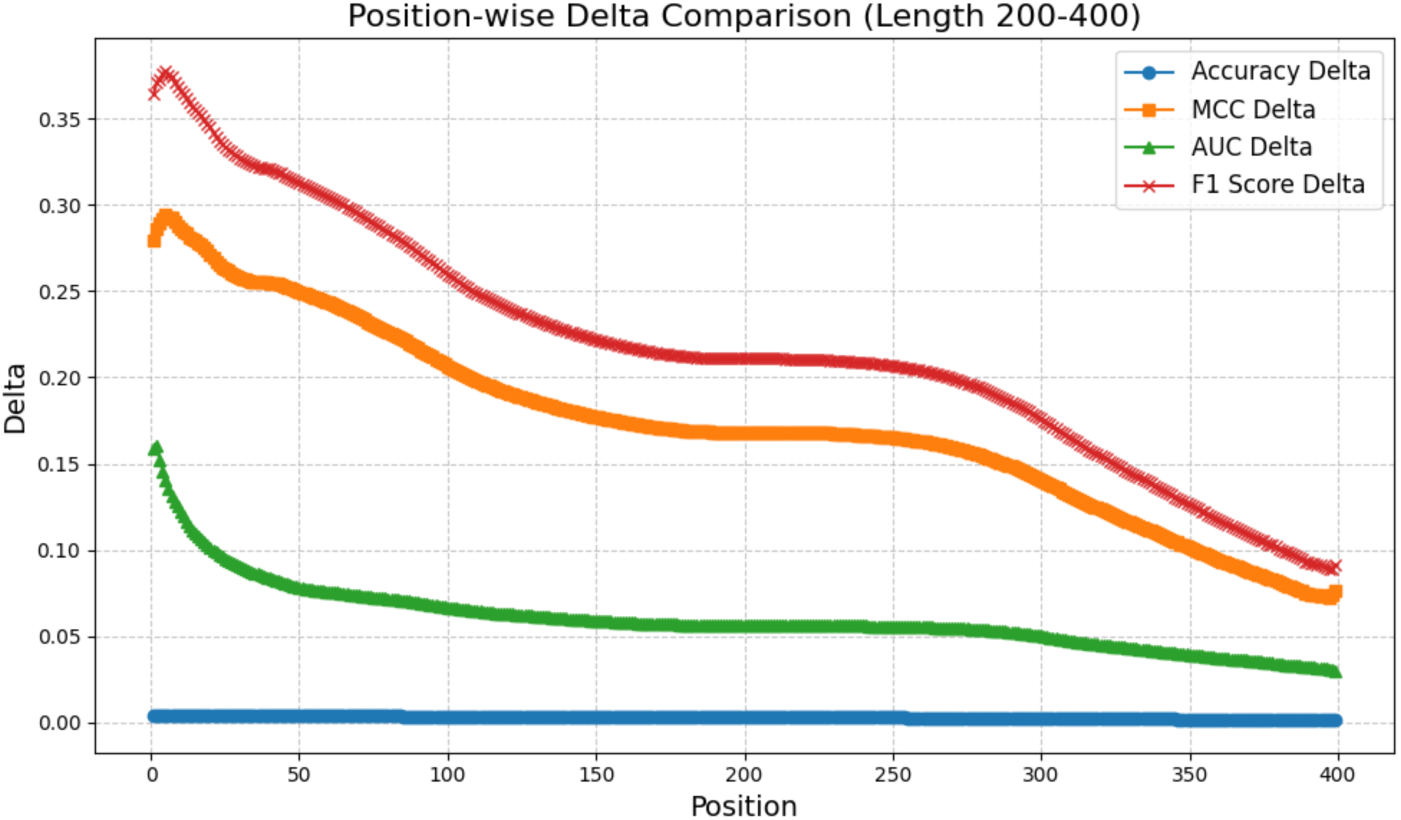
Position-wise delta comparison for sequences of length 200–400. Early positions again dominate the impact on performance, with F1 score deltas exceeding 0.35.

Taken together, these results show a consistent pattern across both length groups: the earlier regions of protein sequences carry the greatest predictive importance for the Lysin-Fusion model, while later positions contribute relatively little. This position-dependent sensitivity highlights the uneven distribution of discriminative features along protein sequences and provides direct evidence that the model relies most strongly on information encoded in the initial segments.

The position-dependent pattern revealed by the occlusion analysis is consistent with the known modular organization of phage lysins. For lysins targeting Gram-positive bacteria, the N-terminal region typically forms the enzymatically active domain (EAD), while the C-terminal region functions as the cell wall-binding domain (CBD)^35^. The binding domain recognizes and attaches to the bacterial cell wall, while the catalytic domain cleaves molecular bonds within the bacterial peptidoglycan^36^. Since our occlusion analysis showed the strongest performance degradation when early positions were masked, it is likely that these positions correspond to the N-terminal EAD, which directly determines catalytic activity and lytic function.

Although the CBD is primarily responsible for anchoring the enzyme to post-lysis cell wall remnants and limiting enzyme diffusion^37^, recent evidence suggests that the EAD also plays a critical role in modulating species selectivity^38^. This indicates that both domains contribute to lysin functionality in distinct ways. Consequently, masking later sequence regions still produced measurable changes in model performance, albeit smaller than those observed at early positions, reflecting the discriminative role of the CBD in lysin classification.

For lysins targeting Gram-negative bacteria, most are single-domain enzymes that lack a CBD^39–41^. These enzymes also carry positively charged groups at the C-terminus that enable penetration or disruption of the outer membrane and facilitate degradation of the peptidoglycan layer^6,42^. However, the catalytic function remains the primary determinant of lytic activity. This structural arrangement is consistent with our observation that occluding early sequence segments produced the largest decreases in model performance, as the functional core resides predominantly in the N-terminal catalytic region.

#### 3.5.2 LIME results

We used LIME to explain the model on the independent test set and aggregated local coefficients across correctly predicted samples. For true positives (TP, lysins), we summed the positive coefficients for class 1; for true negatives (TN, non-lysins), we summed the negative coefficients with respect to class 1 (plotted as absolute values for readability). In both barplots (Figs. 11–12), the *x*–axis lists “feature | interval”, where the interval is the discretized bin used by LIME (for example, “≤ 0.00”, “> 0.01”, or a numeric range); the *y*–axis is the aggregated LIME contribution, and the number atop each bar is its value. Colors indicate encoders: blue = CKSAAP, orange = CTDC, green = APAAC, brown = CTDD.

**Figure 11:**
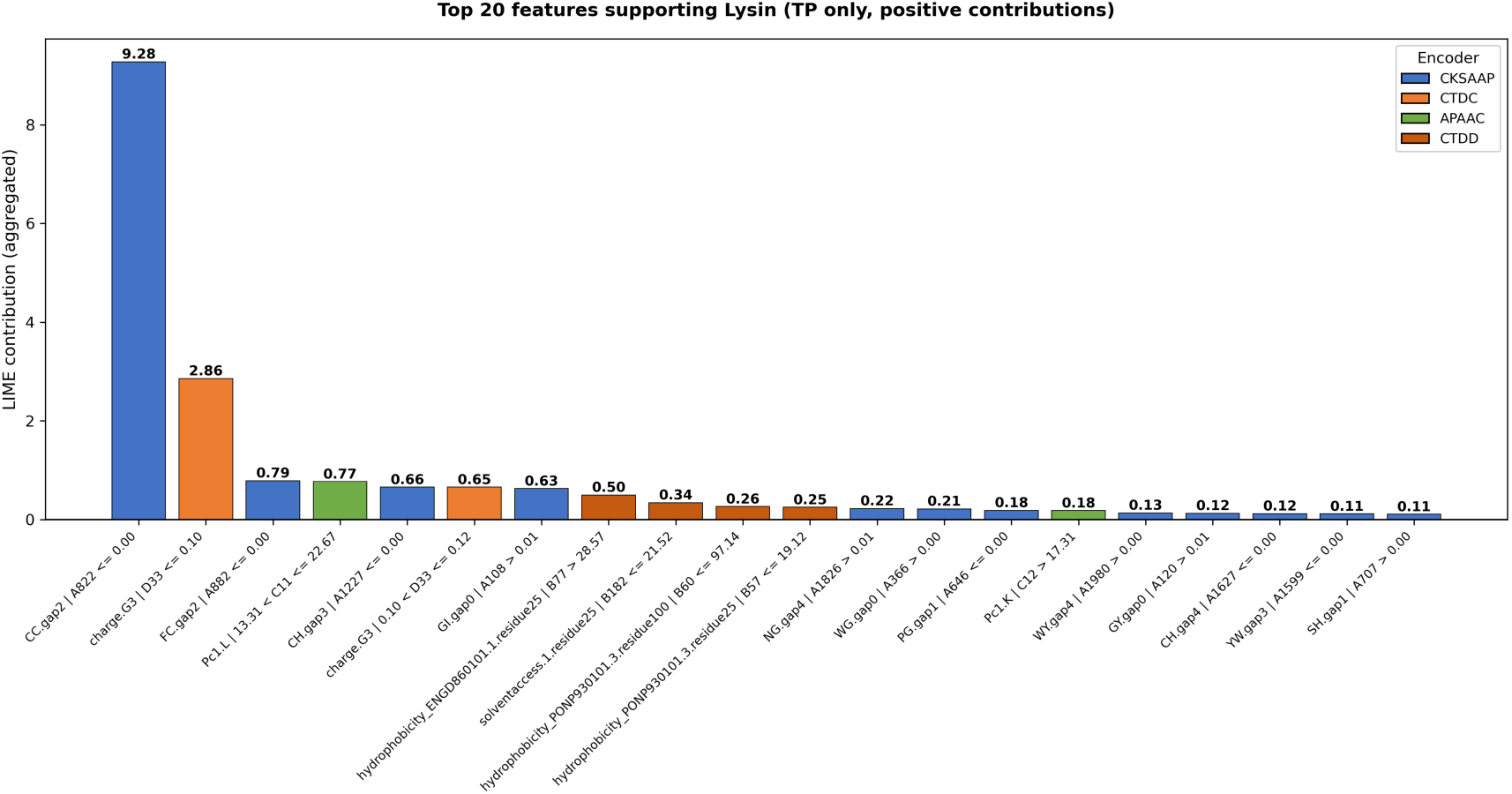
Top 20 features supporting lysin predictions (TP). Each bar shows the aggregated positive LIME contribution. CKSAAP dominates (12 features), with CC.gap2 at the lowest frequency bin showing the largest effect (9.28). CTDC charge composition ranks second (2.86).

**Figure 12:**
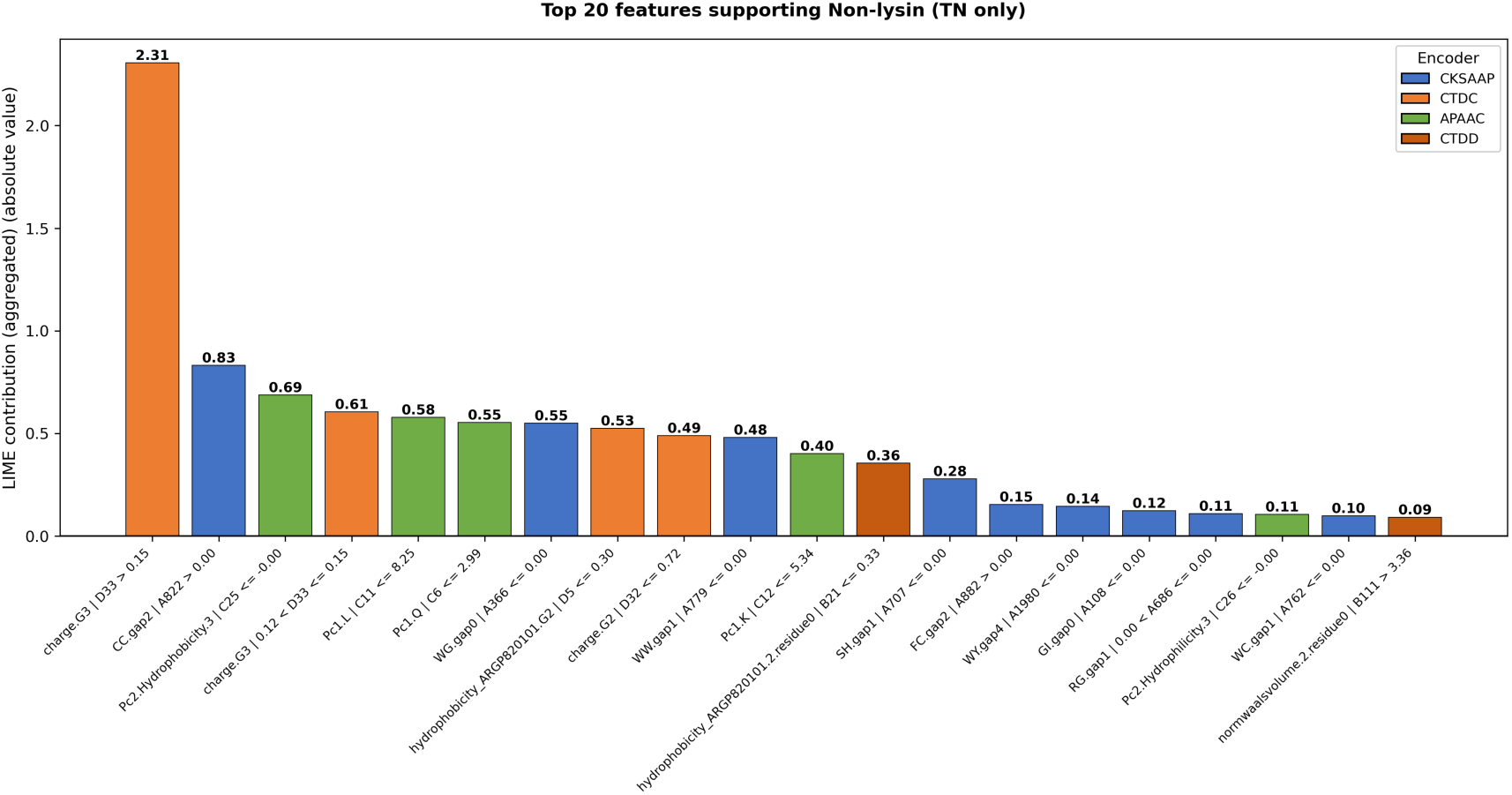
Top 20 features supporting non-lysin predictions (TN). Each bar shows the aggregated absolute value of negative LIME contributions. CTDC charge composition leads (2.31), followed by CKSAAP (0.83). APAAC (5 features) are more prevalent than in the TP top-20.

For lysin-supporting evidence (Fig. 11), CKSAAP provides the dominant signal. The single largest bar is CC.gap2 | A822 ≤ 0.00 with a contribution of **9.28**, showing that when the frequency of the C→C dipeptide at gap 2 falls in the lowest bin (essentially absent), the prediction strongly moves toward lysin. This feature alone accounts for more than three times the contribution of the second-ranked feature. Additional CKSAAP features capture other spaced-dipeptide patterns, such as FC.gap2, CH.gap3, and GI.gap0, most of which also favor lysin when present at low frequencies or in specific value ranges.

The second largest contribution comes from CTDC charge composition: charge.G3 | D33 ≤ 0.10 has a value of **2.86**, indicating that sequences with a low proportion of negatively charged residues (charge group 3) are more likely to be lysins. Another CTDC bin, charge.G3 | 0.10 < D33 ≤ 0.12, also appears in the top 10, reinforcing that this composition threshold is a key decision boundary.

Complementary signals arise from APAAC and CTDD. APAAC features (2 in the top 20) encode sequence-order effects; for example, Pc1.L | 13.31 < C11 ≤ 22.67 (a mid-range leucine composition) contributes 0.77, suggesting that certain amino acid composition ranges combined with their positional patterns support lysin predictions. CTDD features (4 in the top 20) capture where specific property groups occur along the sequence. For instance, hydrophobicity_ENGD860101.1.residue25 | B77 > 28.57 and several hydrophobicity-related quantiles indicate that when certain percentile positions fall into particular bins (e.g., later or earlier along the sequence), the model shifts toward lysin. These APAAC and CTDD contributions are individually smaller (0.1–0.8) but collectively provide fine-grained refinement.

For non-lysin-supporting evidence (Fig. 12), the strongest discriminator is charge composition but in the opposite direction: charge.G3 | D33 > 0.15 has an absolute contribution of **2.31**, and another bin charge.G3 | 0.12 < D33 ≤ 0.15 also ranks high (0.61), both showing that a higher proportion of negatively charged residues supports non-lysins. This directly mirrors the TP result where low charge.G3 supports lysins, confirming that charge composition is a primary decision boundary with complementary thresholds for the two classes.

CKSAAP remains informative but with the complementary interval: CC.gap2 | A822 > 0.00 has a contribution of **0.83**, indicating that when the gap-2 C→C motif is present (frequency above zero), predictions move toward non-lysin. This mirrors the TP finding where its near-absence supports lysin. Overall, CKSAAP accounts for 9 of the top 20 TN features, including patterns like WG.gap0 | A366 ≤ 0.00, WW.gap1 | A779 ≤ 0.00, and SH.gap1 | A707 ≤ 0.00, consistently showing that the presence or absence of specific spaced dipeptides differentiates the two classes.

APAAC features are notably more prominent in the TN top 20 (5 features) than in TP. Low sequence-order signals and low amino acid compositions are associated with non-lysins; for example, Pc2.Hydrophobicity.3 | C25 ≤ 0.00 (0.69), Pc1.L | C11 ≤ 8.25 (0.58), Pc1.Q | C6 ≤ 2.99 (0.55), and Pc1.K | C12 ≤ 5.34 (0.40) all contribute meaningfully, suggesting that weaker or different physicochemical orderings help identify non-lysins.

CTDD features provide positional cues about where property groups occur along the sequence. For instance, hydrophobicity ARGP820101.2.residue0 | B21 ≤ 0.33 and normwaalsvolume.2.residue0 | B111 > 3.36 indicate that the first occurrence of certain hydrophobicity or volume groups tends to fall into particular percentile bins for non-lysins.

Taken together, LIME reveals a clear and interpretable pattern in the model’s decision process. The classifier mainly depends on specific spaced-dipeptide patterns identified by CKSAAP—most notably CC.gap2—whose absence strongly supports lysin prediction while its presence shifts toward non-lysin. Complementary thresholds in charge composition from CTDC (low charge.G3 for lysins, high charge.G3 for non-lysins) further define the class boundary. Finally, fine-grained sequence-order information from APAAC and positional quantiles from CTDD provide additional refinements, contributing more prominently to the discrimination of non-lysins than lysins.

The charge composition patterns revealed by LIME align with documented biochemical characteristics of phage lysins. The identification of charge composition thresholds as a key discriminator is consistent with the critical role of charged residues in lysin function: phage lysins carry positively charged groups at the C-terminus to enable penetration or disruption of the outer membrane and facilitate degradation of the peptidoglycan layer^6,42^. Importantly, LIME identifies lower proportions of negatively charged residues (charge group 3, comprising aspartate and glutamate) as supportive of lysin predictions, which is consistent with the documented enrichment of positively charged residues (lysine and arginine) specifically at the C-terminal region^43,44^. Net charge distribution is significantly higher in Gram-negative lysins than in Gram-positive ones, with this difference concentrated mainly at the C-terminal region and most pronounced at the final sequence quartile^43^. The selectivity of certain lysins has been directly attributed to variations in the C-terminal cationic region^44^. These findings collectively support the critical role of charge composition and distribution in defining the decision boundary between lysins and non-lysins.

The prominence of spaced-dipeptide patterns from CKSAAP, particularly the near-absence of CC.gap2 in lysins, suggests that specific sequence motifs serve as discriminative markers. While the exact functional significance of individual dipeptide frequencies remains to be fully characterized, their strong predictive power indicates that lysins possess characteristic local sequence patterns that distinguish them from other protein families.

The hydrophobicity-related features captured by CTDD correspond to known physicochemical differences. Hydrophobicity is significantly higher in Gram-negative lysins, with Gram-negative lysins exhibiting a more hydrophobic N-terminal region while the C-terminal region shows reversed patterns due to the abundance of positively charged residues. These positional hydrophobicity variations are particularly pronounced at the third quartile, where differences are most statistically significant. Furthermore, hydrophobic moment distributions differ significantly between Gram-positive and Gram-negative phage lysins, with Gram-negative lysins presenting greater hydrophobic moments along most of the protein length except at the N-terminus. These fine-grained positional patterns provide discriminative information that contributes to model performance.

### 3.6 Web server and online prediction system

To facilitate the practical application of our proposed framework, we developed a user-friendly web server for LysinFusion, which provides an accessible interface for lysin prediction based on protein sequences. The web server integrates the complete backend model, including the hybrid CNN–Transformer architecture and multi-feature encoding pipeline, and is freely accessible at: https://huggingface.co/spaces/qq3327008209/LysinFusion

The main interface of the LysinFusion web server is illustrated in Figure 13. The platform supports two prediction modes. For single sequence prediction, users can input the protein name and paste an amino acid sequence into the input box in FASTA format, using only the 20 standard amino acid characters. After clicking the “Run Real Prediction” button, the prediction results will be generated and displayed on a new page.

**Figure 13:**
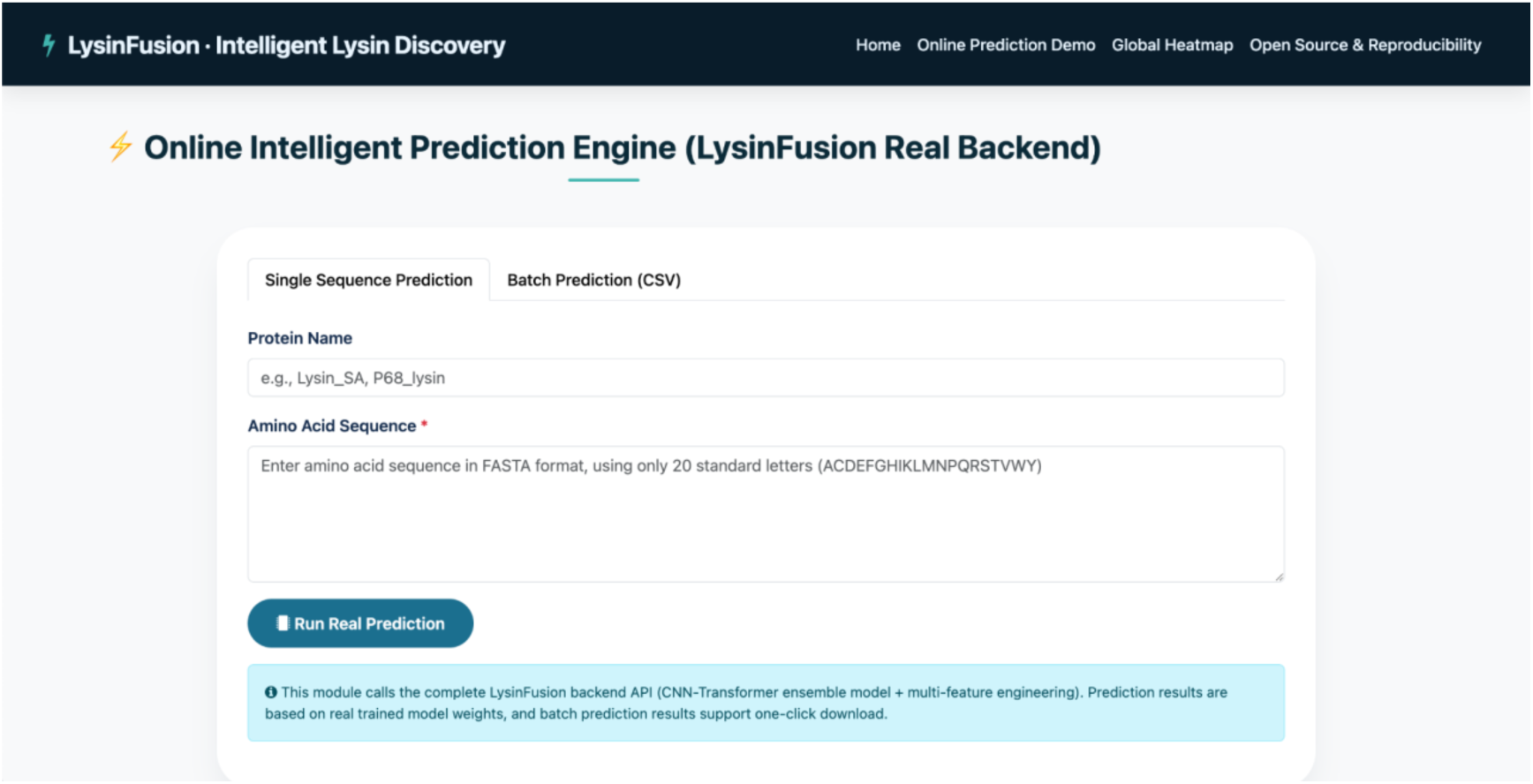
Web interface of the LysinFusion online prediction server. The platform provides two prediction modes: single sequence prediction and batch prediction via CSV upload. Users can input protein sequences in FASTA format and obtain prediction results through the integrated LysinFusion backend model. The interface is directly connected to the trained CNN–Transformer ensemble model with multi-feature encoding, and supports efficient and user-friendly lysin prediction.

For large-scale analysis, the server also supports batch prediction by uploading a CSV file. The uploaded file must contain at least two columns (name and sequence), with the first row specifying the column headers. Batch prediction results can be downloaded with one click after processing.

The web interface is designed to be intuitive and efficient, enabling researchers to quickly obtain prediction results. In addition, the system is directly connected to the trained model weights, ensuring that all predictions are based on the final optimized LysinFusion model. We expect that this web server will provide a convenient and practical tool for high-throughput lysin discovery and related bioinformatics studies.

## 4. Conclusion

We presented LysinFusion, a reproducible framework for binary lysin identification. By combining four complementary sequence encoders (CKSAAP, CTDD, APAAC, CTDC) with a lightweight L1–logistic selector and a hybrid CNN–Transformer classifier, the method jointly captures local motif signals and long-range dependencies. Trained on a curated, de-redundant corpus from PHROG and inphared and evaluated on an independent UniProt benchmark of experimentally validated proteins (148 sequences), LysinFusion consistently outperformed the state-of-the-art DeepMineLys. In particular, it achieved higher overall ACC/AUC/MCC/F1 while markedly reducing false positives, which directly lowers downstream experimental burden. Ablation analyses further confirmed that each major component contributes meaningfully. Importantly, beyond predictive performance, LysinFusion emphasizes reproducibility, interpretability, and practical accessibility through fully available source code, curated benchmark datasets, and an online prediction server, addressing several longstanding limitations of existing lysin prediction tools.

In addition, our interpretability analyses indicate that the model relies on biologically meaningful signals. Occlusion results highlight the early sequence region as most informative, and LIME reveals clear rules: spaced dipeptides from CKSAAP—especially the absence/presence of CC.gap2—and charge composition thresholds from CTDC (low charge.G3 supporting lysins, high charge.G3 supporting non-lysins) delineate primary decision boundaries, while APAAC/CTDD features provide fine-grained refinements. These patterns align closely with known lysin biology: the model’s emphasis on early positions corresponds to the functionally critical N-terminal catalytic domain, while the identified charge composition thresholds reflect the characteristic enrichment of positively charged residues at the C-terminus that enable membrane interaction and peptidoglycan degradation.

This study still has limitations. The training data, although curated and diverse, under-represents rare or highly divergent lysin families, and the current model uses only sequence-level encodings without explicit structural cues. Future work will (i) expand and periodically refresh the benchmark with newly validated lysins and broader taxonomic coverage, including cross-species and large-scale metagenomic screens, and (ii) incorporate structure-aware or pre-trained protein embeddings and pursue model distillation to further enhance generalization and interpretability.

